# A Novel System for The Comprehensive Collection of Nonvolatile Molecules from Human Exhaled Breath

**DOI:** 10.1101/2020.05.14.097113

**Authors:** Dapeng Chen, Wayne A Bryden, Michael McLoughlin

## Abstract

Characterization of nonvolatile molecules in exhaled breath particles can be used for respiratory disease monitoring and diagnosis. Conventional methods for the collection of nonvolatile molecules in breath heavily rely on the physical properties of exhaled breath particles. Strategies taking advantage of their chemical properties have not yet been explored. In the present study, we developed a column system in which the surface chemistry between organic nonvolatile molecules and octadecyl carbon chain was exploited for the comprehensive collection of metabolites, lipids, and proteins. We demonstrated that the collection system had the capture efficiency of 99% and the capacity to capture representative nonvolatile molecules. The collection system was further evaluated using human subjects and proteins collected from human exhaled breath were characterized and identified using gel electrophoresis and bottom-up proteomics. The identified 303proteins from mass spectrometry were further searched against reported bronchoalveolar lavage fluid proteomes and it was shown that 60 proteins have the tissue origin of lower respiratory airways. In summary, we demonstrate that our collection system can collect nonvolatile molecules from human exhaled breath in an efficient and comprehensive manner and has the potential to be used for the study of respiratory diseases.

## Introduction

Human exhaled breath contains nonvolatile molecules that emerge from upper and lower respiratory airways, including small molecules, lipids, and proteins (1–3). Collection of non-volatile molecules in exhaled breath is a non-invasive sampling method and the sequential characterization of those molecules is considered as an attractive strategy for airway monitoring, medical diagnosis, and disease screening (4–6).

The capture of nonvolatile molecules in human breath can be achieved using traditional techniques such as the collection of exhaled breath condensate (EBC) (7). Briefly, EBC collection is accommodated by directing human exhaled breath onto a cooled structure (7). Since there is a temperature gradient between warm, high-humidity exhaled breath air and the cold structure, exhaled air can be condensed and collected as liquid and analyzed. EBC collection methods are advantageous since it only requires tidal breathing and can be feasibly adapted to a non-invasive method for the collection of breath samples from people with compromised lung function (2,7). However, the EBC method has proven to be a very inefficient way to collect nonvolatile molecules from human exhaled breath (8–10). The majority of the particles in exhaled breath are under 1 *μ* m, which are captured with very low efficiency on the cooled surfaces (8,9). In addition, EBC samples contain a large amount of water, which causes a significant dilution of analytes to levels below the detection limits of conventional analytical methods. Considering these drawbacks for EBC methods, alternative methodologies, targeting exhaled breath particles, have been developed as a more aggressive solution. (11–18). Current technologies for collecting exhaled breath particles largely hinge on their size distributions and physical properties (19). Accordingly, exhaled breath particles are often collected using physical impactors using a mouthpiece or cone-like collector with an appropriate flow rate (13–15). Particles contained in the exhaled breath, largely in the form of aerosols, pass through the tubing of the collector and precipitate onto a solid surface. There is a rigorous physical cutoff size of the impactors whereby the small particles are exceedingly difficult to capture (13,14). A combination of filters and impactors is preferred to achieve a more comprehensive collection (18). Technologies relying on the collection of particles provide certain advantages over traditional EBC methods and have been used extensively to investigate lung diseases for biomarker discovery and medical diagnosis (20–23). In addition, the removal of water vapor from exhaled breath largely solves the analyte dilution issues plaguing EBC collection methods (15). Accordingly, nonvolatile molecules captured on solid surfaces can be efficiently collected and prepared for various chemical and biochemical analysis such as immunoassay, polymerase chain reaction (PCR), and mass spectrometry (14, 22, 23).

While current methodologies for the collection of nonvolatile molecules in breath exhaled particles solely rely on their physical properties, strategies taking advantage of their chemical properties have not been fully explored and developed. Nonvolatile molecules in human breath are mostly organic compounds that are less hydrophilic and show a strong affinity for alkyl chains via intermolecular forces including hydrogen bond and noncovalent interaction. These surface chemistry properties have been employed extensively for the development of targeted purification technologies such as solid-phase extraction and reverse phase liquid chromatography (24–27). In this study, we developed a column system in which octadecyl (C18) bonded resin beads were utilized for the comprehensive capture of nonvolatile molecules in human exhaled breath. The system was evaluated in human subjects. Protein samples collected from the exhaled breath were investigated using bottom-up proteomics and the protein profiles were revealed.

## Methods

### Materials

All solvents and chemicals were HPLC-grade and acquired commercially from Fisher Chemical (Waltham, MA). The 10 *μ* m pore-size filter discs and the column sets were acquired commercially (figure 1, Boca Scientific, Dedham, MA). C18 resin beads with 20 *μ* m diameters were purchased from Hamilton (figure 1, Reno, NV).

**Figure 1.**
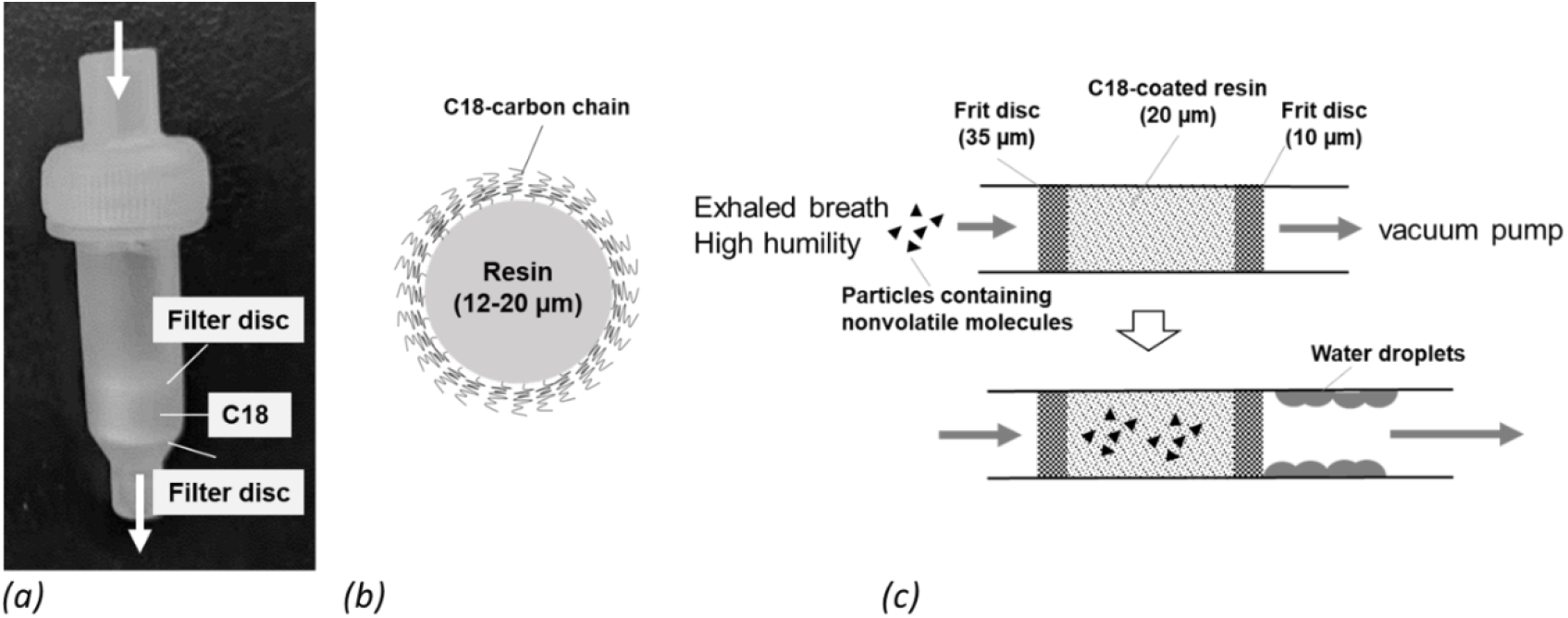
Schematic presentation of the column-based collection system. The packed column system includes two filter discs and C18 resin beads (a, b). Capture mechanism by C18 resin beads from human exhaled breath is described (c).

### Particle capture efficiency testing

The laboratory system for the capture efficiency testing was presented in figure 2a. For the testing, 200 *μ* L of HPLC-grade water was aerosolized into a 2-liter chamber using a portable Aeroneb Go nebulizer (Philips, Amsterdam, Netherlands). Particles counts were recorded using a portable laser particle counter (Met One Instruments, Grants Pass, OR) under 4 experimental conditions: column-only and without C18 (No filter), 0.2 *μ* m pore-size syringe filter with 25 mm ID (VWR International, Radnor, PA), a column equipped with 30 mg of C18 resin beads (C18 column), and the C18 column with 30 min of collection time (30 min incubation).

**Figure 2.**
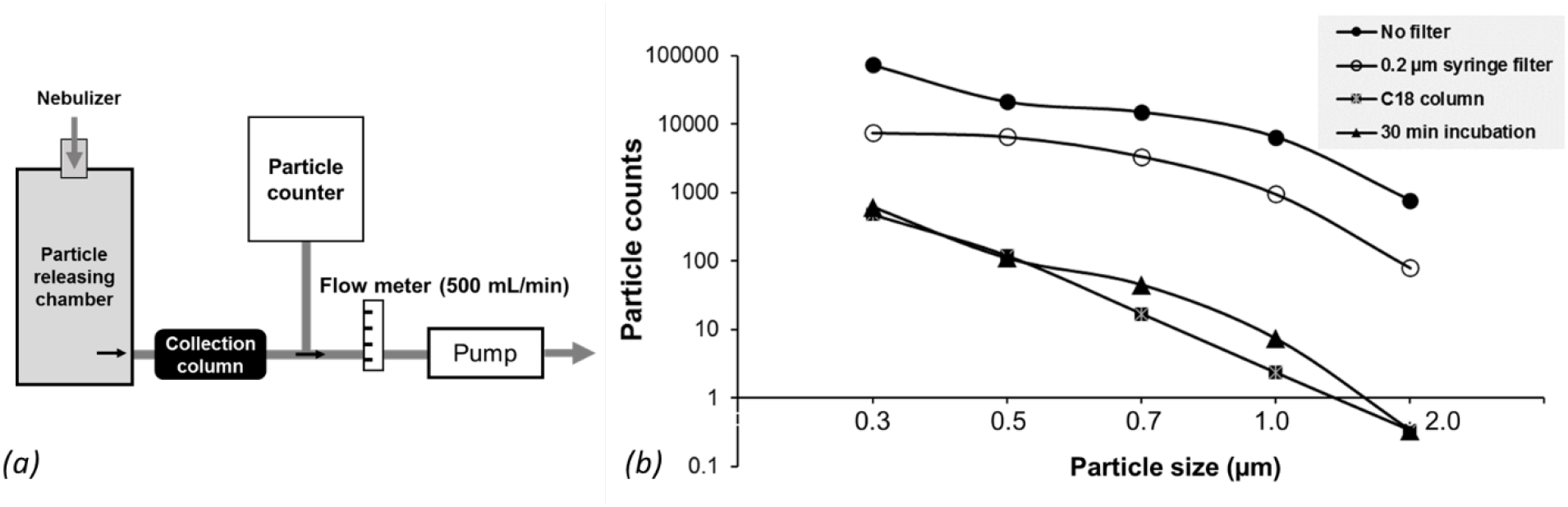
Evaluation of capture efficiency using a laboratory setup. The column collection system includes several components (a) and the results of particle counts under different experimental conditions (b).

### Standard nonvolatile molecule collection testing

The laboratory system for collection capacity testing is presented in figure 3a. The column collection system was evaluated using three representative molecules: methadone (0.001 ng/mL, Sigma-Aldrich, St. Louis, MO), 1,2-Dipalmitoyl-sn-glycero-3-phosphorylcholine (0.1 ng/mL, Matreya LLC, State College, PA), and insulin from porcine pancreas (1 ng/mL, Sigma-Aldrich). Briefly, 200 *μ* L of each molecule was prepared in HPLC-grade water and aerosolized using a portable Aeroneb Go nebulizer (Philips) into a 50 mL conical tube sitting on a heating block (50°C). The collection column was installed on the bottom of the 50 mL conical tube and the flow rate was set as 200 mL/min for 5 minutes. After collection, collection columns used for methadone and insulin were washed with 400 *μ* L of water 4 times. After quick centrifugation, methadone and insulin were eluted using 400 *μ* L of 70% acetonitrile. For the column used for phosphorylcholine collection, the column was washed with 400 *μ* L of 50% acetonitrile 4 times and the elution accomplished using 400 *μ* L of 70% isopropanol. Washing solutions were also collected and saved for analysis.

**Figure 3.**
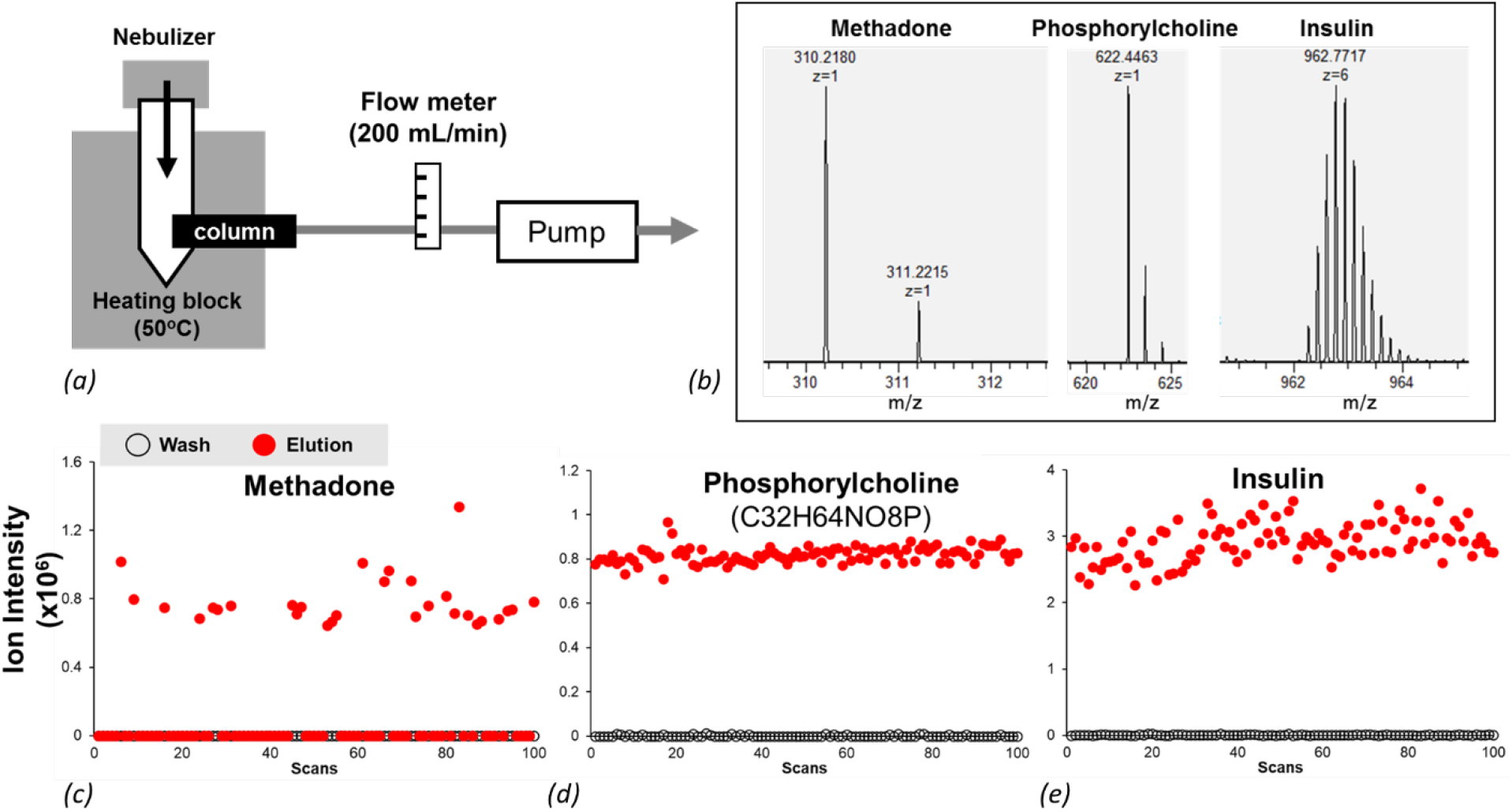
Capture of representative molecules using the collection system. A testing system was built for the molecule releasing and the C18 collection (a). Mass spectra of 3 representative molecules are shown (b) and the ion signals of representative molecules collected from different collection conditions were presented (c-e).

Collected methadone and insulin samples were lyophilized and re-suspended in 200 *μ* L of 70% acetonitrile with 1% acetic acid. The samples containing phosphorylcholine were lyophilized and re-suspended in 200 *μ* L of 50% isopropanol, 25% acetonitrile with 1% acetic acid. Mass spectrometric data collection of methadone, phosphorylcholine, and insulin was conducted on an orbitrap LTQ mass spectrometer via direct infusion in the positive ion mode (Thermo Fisher Scientific). The flow rate of direct infusion was set to 3 *μ* L/min and the data collection was recorded for 10 min at the resolution power of 60,000 at 200 m/z. Identification of the target molecules was conducted by accurate mass measurement.

### Collection of nonvolatile molecules from human exhaled breath

Four healthy volunteers participated in this study. All study participants understood the experimental details and signed the informed consent before the study. This data was analyzed without personal identifiable information and will not be used for regulatory approval. For the collection of nonvolatile molecules from human exhaled breath, a collection column was installed onto a first aid CPR rescue mask with minor modifications (figure 4). To prevent chemical contamination from ambient air particulate, a HEPA filter was connected to the facial mask. Exhaled breath condensate (EBC) collection glassware cooled with an ice water bath was installed after the collection column to evaluate the capture efficiency of the column collection system. The flow rate was controlled by a needle valve and set to with a flowrate of 600 mL/min.

**Figure 4.**
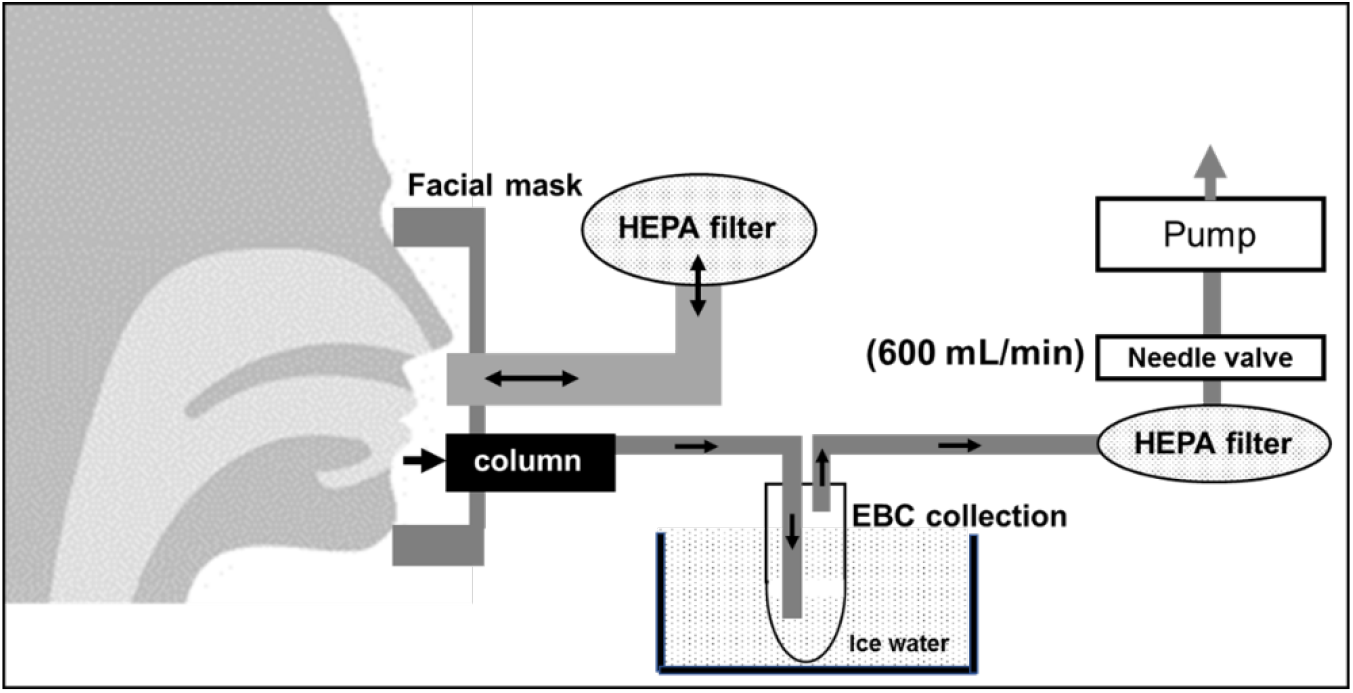
The nonvolatile molecule collection system for human subject study. A HEPA filter and the C18 collection column were installed on a CPR rescue mask. An EBC system was installed to collect molecules that potentially passed through the C18 column. The air flow was assisted by a vacuum pump with the flow rate controlled by a needle valve.

Breath samples were collected from 4 study subjects with different total breathing volume with the same flow rate: subject 1 for 144L, subject 2 for 40 L, subject 3 and subject 4 for 81L. After breath sample collection, the collection column was removed and eluted with 400 *μ* L of 70% acetonitrile for the collection of proteins. The protein samples were lyophilized and resuspended in 100 *μ* L of 0.1% formic acid. Then, the column was eluted with 400 *μ* L of 70% isopropanol for the collection of nonpolar molecules for future analysis.

### SDS-PAGE electrophoresis and silver staining

25 *μ* L of total collected sample was used for SDS-PAGE electrophoresis, which was conducted using a Criterion Tris-HCl Gel system following the manufacture instructions (Bio-Rad Laboratories, Hercules, CA). After SDS-PAGE electrophoresis, the SDS-PAGE gel was prepared with a silver staining kit (Thermo Fisher Scientific) for the visualization of protein bands. Bovine serum albumin (BSA) was used as an internal positive control.

### Bottom-up proteomics

50 *μ* L of total collected sample was used for the bottom-up proteomics. Briefly, 50 *μ* L of 50 mM ammonia bicarbonate (pH 8.5) was added to each sample. Protein reduction was conducted by adding dithiothreitol to a final concentration of 5 mM and incubated for 30 min at 37°C. After reduction, protein alkylation was followed by adding iodoacetamide to a final concentration of 15 mM and incubated for 1 h at room temperature. Trypsin (Thermo Fisher Scientific) was used for an overnight protein digestion. After digestion, peptides were cleaned up using C18-packed tips (Glygen, Columbia, MD). The peptide samples in 20 *μ* L of 0.1% formic acid were then prepared for mass spectrometry analysis.

Samples were processed using an EASY-nLC 1000 system (Thermo Fisher Scientific) coupled to a Q Exactive HF Hybrid Quadrupole-Orbitrap mass spectrometer (Thermo Fisher Scientific). For the tandem mass spectrometry analysis, peptides were loaded into an Acclaim PepMap 100 C18 trap column (0.2 mm × 20 mm, Thermo Fisher Scientific) with a flow rate of 5 *μ* l/min and separated on an EASY-Spray HPLC Column (75 *μ* m × 150 mm, Thermo Fisher Scientific). HPLC gradient was conducted using 5-55% of the mobile phase (75% acetonitrile and 0.1% formic acid) with a flow rate of 300 nL/min for 60 minutes. Mass spectrometry data collection was conducted in the data-dependent acquisition mode. Key parameters were described as following. Precursor scanning resolution was set to 60,000 and product ion scanning resolution 15,000. Product ion fragmentation was accomplished using high energy collision-induced disassociation (HCD) with 27% total energy. The bottom-up proteomics raw data files were processed with MaxQuant-Andromeda software (*maxquant.org*) against the human protein database (*uniprot.org*) following the standard recommendations and instructions.

## Results

The collection system was composed of C18-coated resin beads with two frit discs with an appropriate pose size established on both sides (figure 1a, b). During sample collection, human exhaled breath, composed of ~99% saturated humidity at 37 °C, passes through the column at a constant flow rate assisted by a vacuum pump (figure 1c). Since nonvolatile molecules (dark triangles in figure 1c) contained in the exhaled breath interact with C18 functional groups, these molecules tend to be trapped on the stationary resin beads while hydrophilic molecules, mostly water and aqueous electrolytes in the breath, pass through the column.

A laboratory aerosol particle releasing system, generating particles of 0.3 to 2 *μ* m size, was used for evaluation of the particle capture efficiency of the collection system (Figure 2a). The particle counts for 0.3 *μ* m particles were ~ 37,000 without the C18-packed column and the counts dropped to 480 with the C18-packed column, indicating >99% capture efficiency (figure 2b and table 1). The similar capture efficiency was observed in particles of all sizes (figure 2b and table 1). Most importantly, the high capture efficiency was well preserved after running the collection for 30 minutes (figure 2b and table 1).

**Table 1.** Particle counts of different sizes for the capture efficiency testing.

| Conditions | Particle Sizes |  |  |  |  |
| --- | --- | --- | --- | --- | --- |
| | 0.3 $\mu\text{m}$ | 0.5 $\mu\text{m}$ | 0.7 $\mu\text{m}$ | 1.0 $\mu\text{m}$ | 2.0 $\mu\text{m}$ |
| No Filter | 36592 ±143 | 10703 ±215 | 7747 ±481 | 3434 ±398 | 427 ±91 |
| 0.2 $\mu\text{m}$ syringe filter | 7401 ±2007 | 6485 ±1513 | 3367 ±920 | 950 ±676 | 79 ±64 |
| C18 column | 480 ±131 | 120 ±8 | 17 ±6 | 2 ±2 | 0 ±1 |
| 30 min incubation | 617 ±150 | 110 ±22 | 45 ±9 | 7 ±5 | 0 ±1 |

Based on size and chemical properties, nonvolatile molecules can be divided into three broad categories: small polar molecules, small nonpolar molecules (lipids), and macromolecules (peptides and proteins). To demonstrate that the C18 column collection system has the capacity to capture the molecules of all types, a proof-of-concept study was performed using a laboratory particle nebulizing system modified from figure 2a (figure 3a). Representative molecules from each category were selected and characterized using high resolution mass spectrometry for the accurate mass measurement (figure 3b): methadone represents small polar molecules, phosphorylcholine for lipids, and insulin for peptides and proteins (figure 3b). After collection, the columns were washed completely prior to elution to avoid carry-over contamination. Our results showed that mass spectrometric signals for each representative molecule were only detected in the elution solution (red dots, figure 3c-e) but not in the washing solution (dark open dots, figure 3c-e), suggesting a complete collection of the target molecules by the C18 collection column.

The collection column was then evaluated using human subjects. The exhaled breath was introduced to the collection system using a modified facial mask system (figure 4). To maximize the particle collection, the C18 column was installed in a way that it exposed directly to the exhaled breath but no direct contact between the month and the inlet of the column to avoid saliva contamination (figure 4).

Protein content in human exhaled breath is still in debate due to different techniques employed for the protein content determination. Two main techniques are typically used: colorimetric detection-based protein assay and staining-based gel electrophoresis. Colorimetric detection methods are less technically challenging, but the specificity is often compromised by polymer contaminants during sample collection that cause a false reading, usually much higher than the actual value. On the other hand, gel electrophoresis-based protein band imaging techniques, such as silver staining, are more technically intensive with the trade-off of offering a very high specificity with the visualization of protein molecular masses. In our study, the collected protein samples from human subjects were evaluated using SDS-PAGE silver staining. The intensity of the protein bands suggested the protein content in all samples was over 10 ng, which is adequate for the sequential bottom-up proteomics. In addition, the protein visualization showed a similar protein band pattern among the 4 human subjects as the most intense protein bands were observed in 3 main positions: 50-75 kDa, 37 kDa, and 10-15 kDa (figure 5a). By using tandem mass spectrometry, those proteins bands were identified as serum albumin and keratins at the position of 50-75 kDa (red cross in figure 5a, supplementary table 1), zinc-alpha-2-glycoprotein at 37 kDa (red double-cross in figure 5a), and cystatin, dermcidin, and S100 proteins at 10-15 kDa (red star in figure 5a). In addition, the protein clusters shown at ~37 kDa suggest very heavy protein posttranslational modifications, presumably glycosylation, may occur. Indeed, bottom-up proteomics and protein database searching confirmed that this protein is zinc-alpha-2-glycoprotein, a protein with 4 known N-linked glycosylation sites (table 2 and *uniprot.org*). The results indicate that silver staining-based protein visualization coupled with bottom-up proteomics provides superb advantages to reveal protein profiles in samples with very low protein concentration.

**Table 2.** The top 20 proteins identified from 4 subject samples based on the abundance.

| Accession ID | Protein Identification | Sequence coverage [%] | Mol. weight [kDa] | Sequence length (aa) | MS/MS count |
| --- | --- | --- | --- | --- | --- |
| P01040 | Cystatin-A OS=Homo sapiens OX=9606 GN=CSTA PE=1 SV=1 | 100 | 11 | 98 | 46 |
| P81605 | Dermcidin OS=Homo sapiens OX=9606 GN=DCD PE=1 SV=2 | 43.6 | 11 | 110 | 35 |
| P31151 | Protein S100-A7 OS=Homo sapiens OX=9606 GN=S100A7 PE=1 SV=4 | 88.1 | 11 | 101 | 28 |
| Q9NZT1 | Calmodulin-like protein 5 OS=Homo sapiens OX=9606 GN=CALML5 PE=1 SV=2 | 95.9 | 16 | 146 | 26 |
| P62987 | Ubiquitin-60S ribosomal protein L40 OS=Homo sapiens OX=9606 GN=UBA52 PE=1 SV=2;sp | 57.8 | 15 | 128 | 23 |
| P00441 | Superoxide dismutase [Cu-Zn] OS=Homo sapiens OX=9606 GN=SOD1 PE=1 SV=2 | 99.4 | 16 | 154 | 17 |
| P25311 | Zinc-alpha-2-glycoprotein OS=Homo sapiens OX=9606 GN=AZGP1 PE=1 SV=2 | 41.6 | 34 | 298 | 12 |
| P31944 | Caspase-14 OS=Homo sapiens OX=9606 GN=CASP14 PE=1 SV=2 | 38.8 | 28 | 242 | 11 |
| P01036 | Cystatin-S OS=Homo sapiens OX=9606 GN=CST4 PE=1 SV=3 | 59.6 | 16 | 141 | 11 |
| P02768 | Serum albumin OS=Homo sapiens OX=9606 GN=ALB PE=1 SV=2 | 14.6 | 69 | 609 | 10 |
| P0DP25 | Calmodulin-3 OS=Homo sapiens OX=9606 GN=CALM3 PE=1 SV=1;sp | 38.3 | 17 | 149 | 9 |
| P04264 | Keratin, type II cytoskeletal 1 OS=Homo sapiens OX=9606 GN=KRT1 PE=1 SV=6; | 18.3 | 66 | 644 | 9 |
| Q6UWP8 | Suprabasin OS=Homo sapiens OX=9606 GN=SBSN PE=1 SV=2 | 20.7 | 61 | 590 | 9 |
| P02538 | Keratin, type II cytoskeletal 6A OS=Homo sapiens OX=9606 GN=KRT6A PE=1 SV=3; | 15.1 | 60 | 564 | 7 |
| P13645 | Keratin, type I cytoskeletal 10 OS=Homo sapiens OX=9606 GN=KRT10 PE=1 SV=6; | 7.4 | 59 | 584 | 6 |
| P31025 | Lipocalin-1 OS=Homo sapiens OX=9606 GN=LCN1 PE=1 SV=1;sp | 19.3 | 19 | 176 | 5 |
| O14529 | Homeobox protein cut-like 2 OS=Homo sapiens OX=9606 GN=CUX2 PE=1 SV=4 | 1.9 | 162 | 1486 | 4 |
| Q6E0U4 | Dermokine OS=Homo sapiens OX=9606 GN=DMKN PE=1 SV=3; | 10.5 | 47 | 476 | 4 |
| Q08554 | Desmocollin-1 OS=Homo sapiens OX=9606 GN=DSC1 PE=1 SV=2 | 5 | 100 | 894 | 4 |
| Q02413 | Desmoglein-1 OS=Homo sapiens OX=9606 GN=DSG1 PE=1 SV=2 | 6 | 114 | 1049 | 4 |

**Figure 5.**
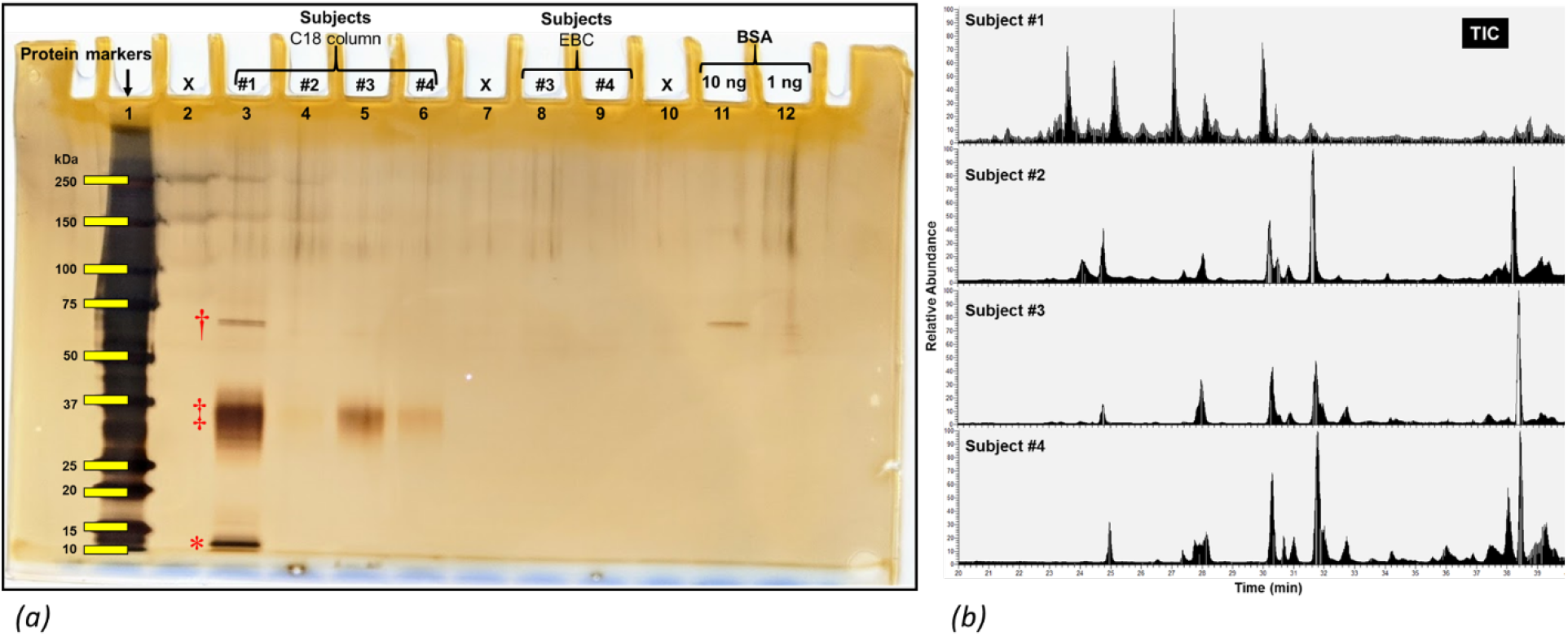
Characterization of proteins collected from human breath samples using silver staining and lipid chromatography mass spectrmetry. Silver staining images with SDS-PAGE electrophoresis was used to visualize protein bands from different human subjects (a). Peptide profiles produced from proteins collected from 4 human subjects were characterized using total ion chromatography (TIC) with mass spectrometry (b).

Another important observation is that the intensity of the protein bands is consistent with the collected breath volume as subject 1, being collected for 144 liters of breath, showed the darkest protein bands as subject 2 showed the lightest bands (figure 5a). The similarity in protein pattern was further supported by the total ion chromatography (TIC) in LC-MS analysis (figure 5b). The major ion chromatography peaks among the 4 subjects were overlapped as subject 1 showed more intense peaks due to the longer sample collection time (figure 5b).

Bottom-up proteomics was used for the identification of proteins contained in the collected exhaled breath samples. 197 proteins were identified from subject 1, 47 proteins from subject 2, 25 proteins from subject 3, and 64 proteins for subject 4 (supplementary table 1). The protein identification numbers are consistent with the protein content in the samples as subject 1 had the most proteins identified. In total, 303 proteins were identified from the 4 subjects. The most abundant proteins identified based on spectral matching were listed in Table 2, including cystatin-A, dermcidin, and several members in the S100 protein family.

As previously mentioned, nonvolatile molecules contained in the exhaled air could be from both upper and lower respiratory airways. To reveal the tissue origin of the proteins identified in our study, we studied 5 published proteome databases from bronchoalveolar lavage fluid (BALF) studies as the BALF sampling method represents the exclusive origin of lower respiratory airways (supplementary table 2) (28–32). The comparison results showed that 63 proteins identified in our study were reported in the BALF proteomics, suggesting our collection system was able to capture proteins shed from lower respiratory airways (table 3).

**Table 3.**
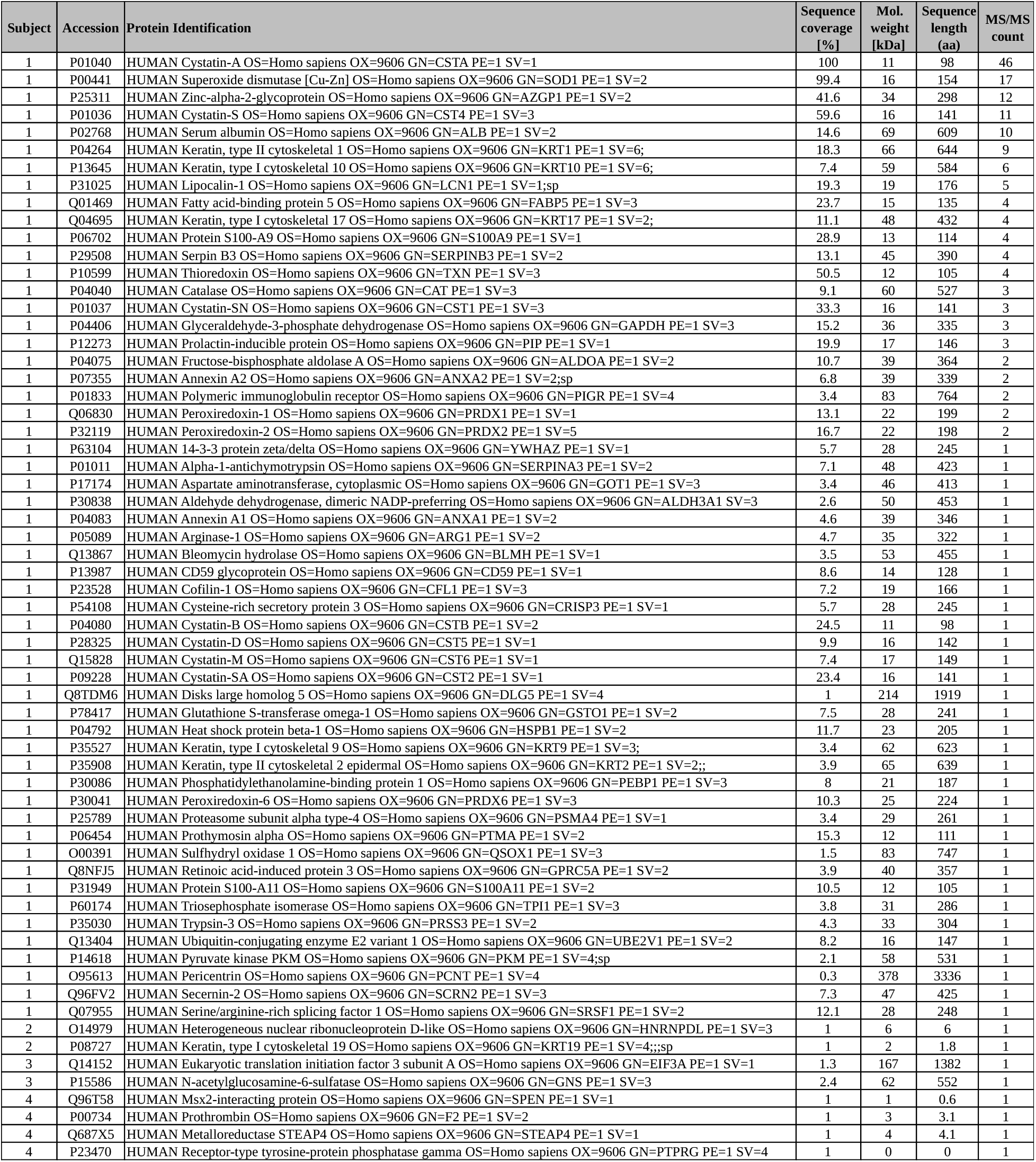
Proteins list of identifications from both the current study and BALF proteomes.

## Discussion

In the current study, we presented a C18 column-based system for the collection of nonvolatile molecules from human exhaled breath. We demonstrated that this collection system had high efficiency and comprehensive capacity to collect particles of all sizes and all types. Moreover, we applied the collection system to human subjects and evaluated the proteins identified from breath samples using published BALF proteome databases. The identification of lower respiratory airway proteins in our study suggests that the collection system has the potentials to be used for a diagnostic tool based on protein profiles.

It is well known that aerosol particles containing nonvolatile molecules have a broad size distribution ranging from sub-micron to several microns, which requires that a collection system should be sophisticated enough to capture particles of all sizes (19, 33). Accordingly, our initial testing focused on capture efficiency and we demonstrated that the C18-based collection system achieved >99% capture efficiency for all particle sizes. Most importantly, the capture efficiency was well preserved after a long term of sample collection, which is quite important for the concentration of target molecules to improve the detection level. Besides having a broad particle size distribution, breath aerosol particles contain a variety of organic biomolecules such as metabolites, lipids, and proteins, which means that a collection system should hold the capacity to capture organic and biological molecules of all types. The C18 resin material that we chose for our study has been widely used in reverse phase chromatography and solid-phase extraction in metabolomics, lipidomics, and proteomics, which indicates that this material should capture organic molecules in their aerosol form (24). Indeed, in the proof-of-concept study, we demonstrated that the collection system had the capacity for the complete capture of three representative molecules in a very comprehensive fashion.

After our initial testing, we tested the collection system to evaluate whether it would be sophisticated enough to capture nonvolatile molecules contained in human breath. There is no clear evidence whether, or at least in what proportion, human breath particles contain nonvolatile molecules from the lower respiratory airways deep in the lungs. Analysis of protein profiles serves this purpose well since proteomics data is an excellent method to reveal the tissue origin. Accordingly, we implemented two strategies for a better understanding of the tissue origin of the protein identified collected from exhaled breath samples. The initial analysis focused on whether the column system has the capacity to capture proteins contained in exhaled breath. For this purpose, we took advantaged a gold standard protein imaging technique, SDS-PAGE silver staining. This system avoids false positive results commonly observed from colorimetric reaction-based protein detection methods due to external contaminants such as detergents, polymer materials, and organic solvents (private conversations with experts in protein chemistry and in lab meetings). The protein bands from silver staining were visualized in all the human subject samples, indicating the protein content in the collected breath samples went beyond the detection level of silver staining, >15 ng *per* protein band. Moreover, similar protein band patterns were observed among the samples with differences in intensity that correlated well with the sample volume collected. In addition, it was reported that different breathing manners affected aerosol particle output, which could contribute to an uneven protein distribution in different subjects as there was no specific instruction given on the breathing manner in this study (34). The similar protein pattern was further confirmed in the TIC profile using tandem mass spectrometry. By using different sophisticated protein analysis techniques, the results strongly suggested that the column collection system was able to capture proteins from exhaled breath and the protein content in the breath samples supported sequential protein identification using bottom-up proteomics.

Bronchoalveolar lavage fluid (BALF) is a sampling method that has been used extensively to investigate pathogenic infections of the lower respiratory system (28–32). Accordingly, the molecular signatures collected from BALF analysis represent an exclusive tissue origin of the lower respiratory airways. Therefore, we investigated various BALF proteomes from previously reported studies to reveal the potential tissue origin of proteins identified in our study. Regardless of different techniques employed for BALF sample preparation and analysis, it can be concluded that several groups of proteins were unambiguously identified in BALF proteomes, including human serum high-abundant proteins such as albumin, S100 protein family, and glycoproteins in lipid metabolism (28–32). The comparison revealed that around 21% of proteins identified in this study overlapped with reported BALF proteomes, strongly suggesting the proteins collected using the column system from exhaled breath samples have the tissue origin of lower respiratory airways. This notion is further confirmed by the observation that the most abundant proteins in this study overlapped with BALF proteomes, including cystatin-A, superoxide dismutase [Cu-Zn], zinc-alpha-2-glycoprotein, cystatin-S, serum albumin, and protein S100-A9 (28–32). Previous studies demonstrated that saliva would be the most likely contamination source for breath sample analysis (2). This does not seem to be the case in our study. First, the modifications of the facial mask (figure 4) avoided direct contact with the month. Moreover, we investigated the saliva protein profiles using silver staining and overlap of the major protein bands between the saliva and breath samples were not observed (supplementary figure 1). In addition, major proteins in saliva, such as alpha amylase, were not identified in our study (35). Based on these observations, it is suggested that our collection system presented in this study can capture proteins from the lower respiratory airways.

In conclusion, we presented a column collection system that can effectively and comprehensively capture nonvolatile molecules in the form of aerosol particles. This collection system was further applied to human subjects and the results demonstrated its capacity to collect proteins from the lower respiratory airways. It is suggested that the presented collection system can be employed to study molecular signatures of lung diseases for early monitoring, diagnosis, and biomarker discovery. Furthermore, this column system has the potential to be developed to capture and study pathogens in both upper and lower respiratory airways.

## Supporting information

STable02

STable01

## Acknowledgements

We thank Tom McCreery (Zeteo Tech, Inc) and Dr. Charles Call (BioFlyte, Inc) for the feedback on the manuscript.

## Author contributions

D.C, M.M, and W.A.B designed the experiments. D.C. collected breath samples. D.C. conducted mass spectrometry analysis and other analysis. D.C. drafted the main manuscript. All authors understood and agreed the results. All authors approved the manuscript.

## Data availability

Data presented in this study are included in this paper. Mass spectrometry data can be communicated with the corresponding author.

## Supplementary materials

Fig. S1. Silver staining images of human exhaled breath and human saliva samples.

Table S1: Protein identifications from 4 human breath samples with the collection system.

Table S2: Reported proteomes of bronchoalveolar kavage fluid from previous studies.

**Supplementary figure 1.**
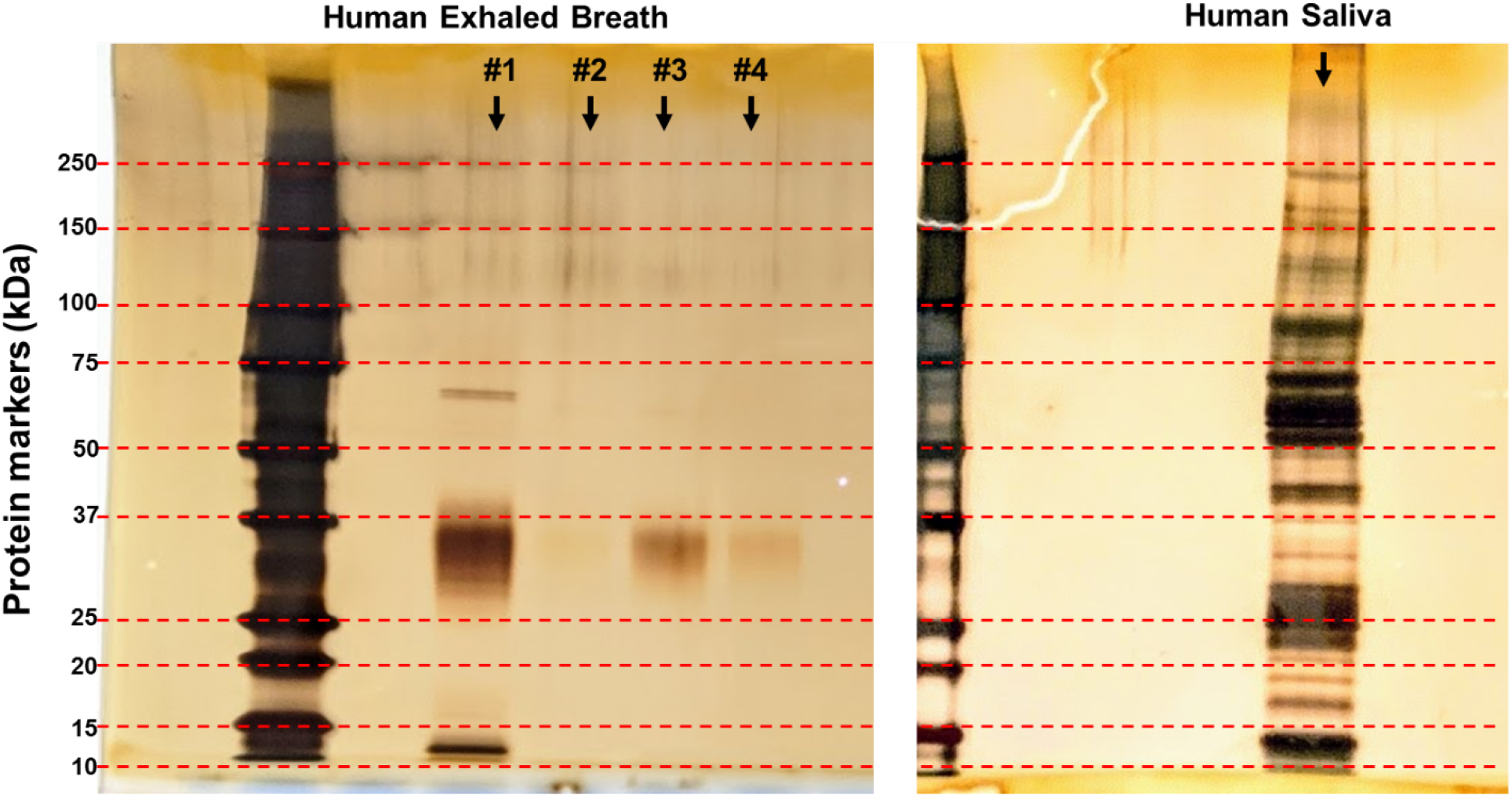
Protein comparison between breath and saliva samples.

